# Preventing Disease Emergence Following Eradication: Application to Mpox

**DOI:** 10.64898/2026.03.03.709287

**Authors:** Cora Hirst, Julia Deichmann, Ananya Saha, Ira Longini, Andreas Handel, Marc Lipsitch, Daniel Weissman, Rustom Antia

**Affiliations:** Department of Biology, Emory University, Atlanta, GA, USA; Department of Epidemiology, Harvard T.H. Chan School of Public Health, Boston, MA, USA; Center for Communicable Disease Dynamics, Harvard T.H. Chan School of Public Health, Boston, MA, USA; Department of Biostatistics, University of Florida, Gainesville, FL, USA; Department of Epidemiology and Biostatistics, University of Georgia, Athens, GA, USA; Division of Infectious Diseases and Geographic Medicine, Department of Medicine, Stanford University, Stanford, CA, USA; Department of Biology, Stanford University, Stanford, CA, USA; Center for International Security and Cooperation, Freeman-Spogli Institute, Stanford University, Stanford, CA, USA; Department of Physics, Emory University, Atlanta, GA, USA

## Abstract

We use simple mathematical models to explore the factors that influence the evolutionary emergence of mpox to a pathogen capable of sustained human to human transmission that poses a global threat. Smallpox eradication followed by the discontinuation of immunization with vaccinia has led to a decline in the level of population immunity against related poxviruses such as mpox. This decline in immunity results in an increase in both the number of spillovers and the extent of human to human transmission. We find that increases in transmissibility of mpox between humans have a much greater effect on the probability of evolutionary emergence compared with increases in the number of zoonotic spillovers. We suggest that while mpox only needs to have a reproductive number slightly greater than one to become endemic, subsequent adaptation is likely to further increase its transmissibility in the human population. As a consequence a much higher level of vaccination (or other intervention) is needed to control the pathogen after its evolutionary emergence compared with what is needed to prevent it from emerging in the first place.

## 1 Introduction

Our understanding of the use of vaccines and other intervention measures for the control and eradication of highly transmissible infectious pathogens was facilitated by the insights provide by relatively simple mathematical models of disease transmission and control [3, 27]. These models predict how the prevalence of a disease depends its basic reproductive number *R*_0_ which equals the number of secondary infections arising from an infected individual introduced into a completely susceptible population [19]. Disease eradication requires vaccine coverage to exceed a fraction 1 − 1*/R*_0_ of the population. The successful implementation of vaccination strategies inspired by consideration of the above heuristic led to the global eradication of two diseases: smallpox in humans (1980) [5] and rinderpest in cattle (2011) [28]. Because smallpox lacks a zoonotic reservoir (humans are the sole hosts for virus), vaccination against smallpox was discontinued following eradication [7, 11, 35].

While there is no zoonotic reservoir in which smallpox circulates, related orthopoxviruses, which we generically refer to as mpox, do circulate in a number of mammalian hosts and have the ability to infect humans [7, 15]. Historically, mpox did not pose a threat to human populations as there were relatively few zoonotic cases and very limited human-human spread from these spillovers [8]. This arose because immunity to smallpox at the population level (herd immunity) conferred cross-reactive protection against mpox [15]. The cessation of mass vaccination against smallpox and the consequent decline in the level of population immunity has potentially opened a niche for the emergence of mpox [6, 24, 32, 33].

In this paper we develop a simple framework analogous to that used for disease eradication to explore how waning herd immunity to smallpox can facilitate the adaptation of mpox to humans and its emergence as a highly infectious disease. We use this framework to understand how the probability of evolutionary emergence of mpox increases over time, and how interventions can prevent or at least substantially delay the process of evolutionary emergence. Our goal is to develop the framework required for understanding of the key factors that govern the evolutionary emergence of zoonoses following eradication of human pathogens. We then bring our models into contact with the empirical observations of the spillover and transmission of mpox infections in the human population in the past decade, and highlight knowledge gaps that prevent us making accurate quantitative predictions for the evolutionary emergence of mpox.

## Results and Discussion

We clarify our use of reproductive numbers. *R*_0_, the basic reproductive number of a pathogen equals the typical number of secondary cases arising from introduction of an infected individual into a wholly susceptible population. Evolutionary changes resulting in adaptation of a pathogen to humans will result in an increase in *R*_0_. The effective reproductive number *R*_*e*_ of the pathogen depends on its *R*_0_ as well as the extent of immunity in the population. Thus adaptation leads to an increase in *R*_0_ and *R*_*e*_ while a decrease in population immunity leads to an increase in *R*_*e*_ without affecting *R*_0_.

### Outline of the process of emergence

In Fig 1 we outline the stages for the emergence of a novel pathogen [30]. The first stage is cross-species transmission, termed spillover, that results in the introduction of infections into the human population. These spillover events can lead to chains of transmission of the pathogen in the population. If the effective reproductive number *R*_*e*_ *<* 1, then these chains of transmission will stutter to extinction. Evolution of the pathogen during these stuttering chains of transmission can result in an increase in both *R*_0_ and *R*_*e*_ and, if the resulting *R*_*e*_ *>* 1, this can allow mpox to persist in the human population [6, 16]. If *R*_*e*_ has increased to above 1 prior to mpox evolving then further evolution is not needed for it to persist in the human population [42]. In addition to the these stages (which have been considered in earlier models) we introduce a final stage which involves the further adaptation of mpox for transmission in the human population. We expect this final stage to end when the pathogen has reached a fitness peak for transmission in the human population, after which further adaptive evolution will stop or be greatly reduced – for mpox, this might occur when the evolved mpox variant has a basic reproductive number close to that of smallpox.

**Figure 1:**
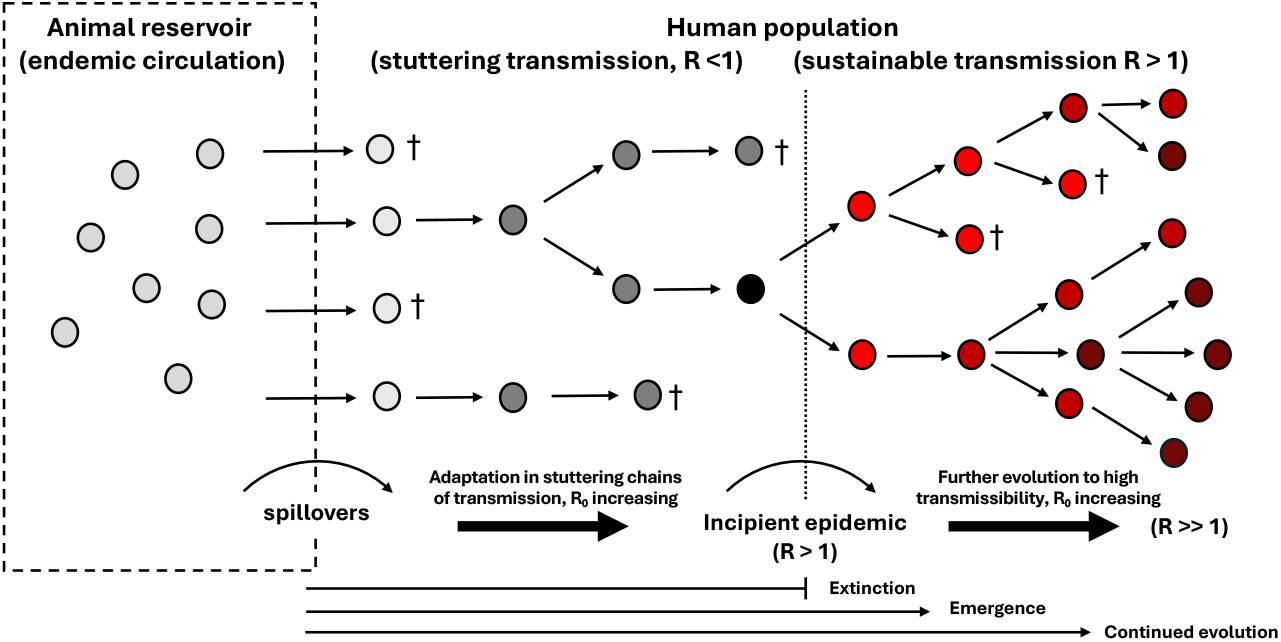
Schematic showing the stages for the emergence of a pathogen. Initially the pathogen causes infections the zoonotic reservoir (circles) and these occasionally spill over into the human population. If the initial reproductive number, *R*_*e*_, of the pathogen in humans is less than one (grey circles) it can nevertheless cause chains of transmission that stutter to extinction. During these chains of transmission *R* can increase (depicted by darker color) and if *R* becomes greater than one (red circles) it can persist in the human population. Adaptive evolution can increase *R* further ending when it has reached a fitness peak in humans (darker red circles).

Applying this framework to mpox requires considering how the progress through these stages is affected by the waning of population immunity following the cessation of vaccination against smallpox.

### Waning immunity and the reproductive number of mpox

We now consider the relationship between the waning of population immunity and the concomitant increase in the *R*_*e*_ of mpox. At least three factors affect the waning of immunity to mpox following smallpox eradication and the cessation of vaccinia immunization [37]. The first is the decrease in the fraction of the population who are old enough to have been immunized against smallpox. The second factor is the waning in the level of immunity to smallpox over time in vaccinated individuals. The third factor is the extent to which smallpox immunity provides immunity to mpox and other orthopoxviruses. Of these factors only the first can be reasonably accurately estimated. Factors that complicate estimation of the level of smallpox immunity include: variation in the magnitude of immunity due to individuals receiving multiple (and different numbers) doses of the vaccine doses; and uncertainties in our quantitative knowledge of how different components of immune memory such as antibodies, CD4 and CD8 T cells wane over time [1, 2, 34] and the relationship between the level of smallpox immunity and protection against mpox. The extent smallpox immunity provides protection against mpox is not only hard to measure but will also depend on the mpox or orthopoxvirus of concern. Finally vaccine efficacy is usually measured as a reduction in number of severe infections while the relevant measure for the emergence of mpox is a reduction in transmission of the virus. We now describe a simple model that gives us a picture of how the transmission of mpox changes over time as the level of population immunity to smallpox declines and the reader is referred to Taube et al [37] for a more detailed consideration.

If we assume that immunity to smallpox is lifelong and all individuals were immune to smallpox when vaccination ended, then we expect population level immunity to smallpox *F* (*t*) to decline as the fraction of the population born prior to the time when smallpox vaccination ceased (circa 1980):

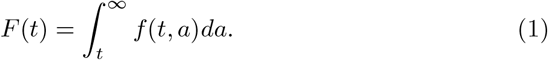

where *f* (*t, a*) is the fraction of population of age *a* at time *t* after smallpox vaccination was stopped. We use estimates for the demographics of the Democratic Republic of Congo (DRC) to get an idea of how the frequency of individuals with smallpox immunity declines and the reproductive number of mpox increases over time [39], and this is shown by the black line in Fig 2.

**Figure 2:**
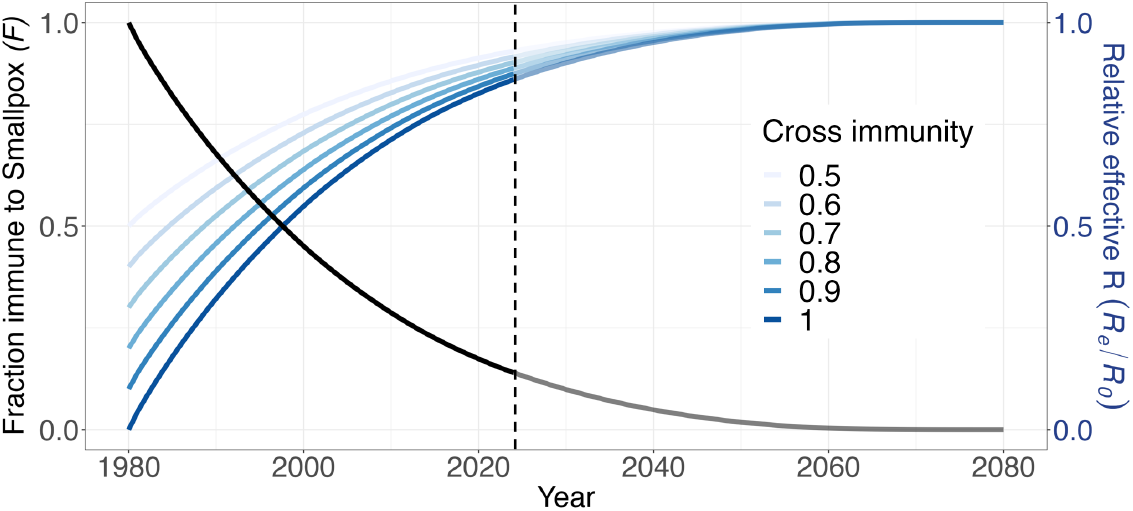
Waning smallpox immunity and the effective reproductive number scaled to its value in a naive population (i.e. *R*_*e*_*/R*_0_) for mpox. The left y-axis and black line indicates how the fraction of individuals in the DRC with immunity to smallpox wanes after vaccination was stopped in 1975. The right y-axis and blue lines indicate how the effective reproductive number of mpox transmission in the human population increases over time. Forward projections after 2023 (vertical dashed line) shown in lighter colors. We assume the fraction of the population immune to smallpox to equal the fraction of individuals born prior to 1980 (the date around which smallpox vaccination was stopped).

The effective reproductive number of mpox, *R*_*e*_ depends both on the extent of immunity to smallpox, *F* (*t*) and the extent to which this provides protection against mpox which we describe by the parameter *c*:

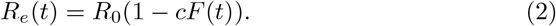

It is clear that the *R*_*e*_ of mpox was much less than one at the time of eradication of smallpox around 1980. Early contact tracing studies suggested *R*_0_ of mpox was slightly less than 1 [12]. The length of stuttering chains of transmission of mpox outbreaks in humans between 1980 and 1984 [9] suggest that *R*_*e*_ at this time was about 0.3. Using estimates of *c* ~ 0.85 [20, 41] and *F* ~ 0.9 at this time, eqn 2 suggests that *R*_0_ is in the range of 1. That the *R*_0_ of mpox is in the range of 1 and likely slightly greater than one is congruent with the relatively slowly growing outbreaks of clade 1 and 2 variants that are currently contained as we consider in the Discussion section.

The broad picture, illustrated in Fig 2 indicates an initially relatively rapid increase in the *R*_*e*_ of mpox from a value much less than 1 at the time of cessation of smallpox vaccination. The rate of increase declines over time reaching a final value that equals the *R*_0_ of mpox, which as indicated above is likely in the region of 1. Because a large fraction of immunity has been lost at the population level, the current *R*_*e*_ of mpox is likely very close to its *R*_0_, and this holds for a large range of potential cross-protection conferred by smallpox immunity (Fig 2).

### Relationship between *R*_*e*_ and evolutionary emergence

The change in population immunity to mpox affects its emergence by increasing the number of spillovers and also the number of secondary cases arising from each spillover. We consider the effect of these two factors on the emergence of mpox.

#### The rate of spillovers

The rate of spillovers (i.e. the number of spillover infections per unit time) *S*(*t*) equals the number of individuals who would be exposed to and infected following contact with the zoonotic reservoir in the absence of population immunity to smallpox (*S*_0_) and the extent of protection provided by immunity to smallpox. The fraction of individuals with smallpox immunity is described by *F* (*t*) and the level of protection smallpox immunity provides against mpox is *c*, which gives us:

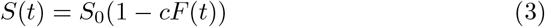

#### Probability of evolutionary emergence per introduction (*P*_*E*_)

We define the probability of evolutionary emergence, *P*_*E*_ as the probability a single spillover results in the emergence of a highly transmissible variant of the pathogen. We use a branching process model of disease transmission (adapted from [4, 6]) to consider how *P*_*E*_ depends on *R*_*e*_, the reproductive number of the introduced virus. In this approach the introduction of an infection from the zoonotic reservoir is followed by stochastic chains of transmission with variable numbers of subsequent infections. If *R*_*e*_ *<* 1, in the absence of pathogen adaptation, these chains of transmission will stutter to extinction. Adaptive evolutionary changes (termed mutations) in the virus can however result in an increase in its reproductive number, *R*_0_ (and consequently *R*_*e*_). The probability of emergence of a zoonotic infection from *R*_*e*_ *<* 1 (subcritical) to *R*_*e*_ *>* 1 (supercritical) depends on the reproductive number of the initial zoonotic infection (*R*_*e*_), the beneficial mutation rate per infection (*µ*) (which we subsequently call simply the mutation rate) and the fitness advantage conferred by a mutation (*σ*), which we assume to be constant (i.e., we assume that there is no macroscropic epistasis [17]). We incorporate two additional features to the earlier model. First, as the effective reproductive number *R*_*e*_ of mpox increases fewer adaptive mutations are required for the reproductive number of the evolved variant to exceed one. Second, we allow adaptive evolution to continue after emergence. We let this occur until the reproductive number of mpox reaches a value comparable to that of smallpox (i.e. *R*_0_ = 3), at which point the epidemic will be much harder to control. We refer the reader to the supplement for the branching process calculations.

In Fig 3 we plot how *P*_*E*_ depends on *R*_*e*_, *µ*, and *σ*. The plots illustrate how *P*_*E*_ increases with increasing *R*_*e*_. If *R*_*e*_ *>* 1, *P*_*E*_ is relatively high, and only stochastic extinction when the numbers of infecteds is low prevents emergence. Fig 3A shows that *P*_*E*_ can fall very rapidly as *R*_*e*_ decreases below 1, and the dependence of *P*_*E*_ on *R*_*e*_ is stronger when the benefit arising from each mutation (*σ*) is small. This is because as *σ* decreases the number of mutations (*n*) required for *R* to exceed one increases as approximately (1 − *R*_*e*_)*/σ*. Fig 3B shows how *P*_*E*_ declines as the mutation rate *µ* falls. When *R*_*e*_ *<* 1 the dependence of *P*_*E*_ on *µ* is compounded by the number of mutations required in order for *R* to exceed one as ≈ *µ*^*n*^.

**Figure 3:**
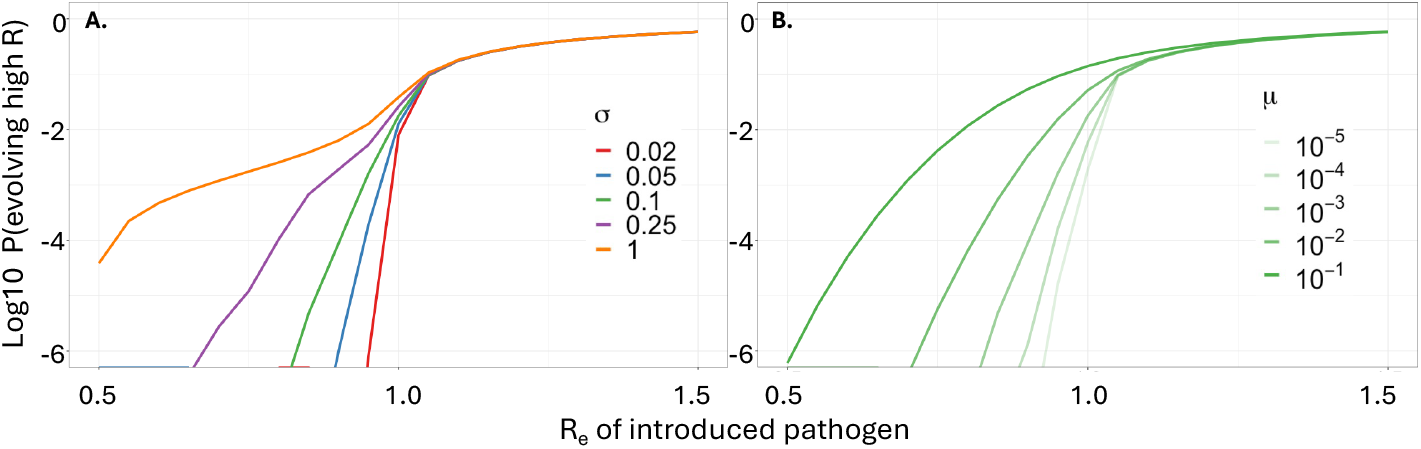
*P*_*E*_ increases with *R*_*e*_, *σ*, and *µ*. The probability of evolutionary emergence per introduction (*P*_*E*_) for a pathogen depends on the effective reproductive number *R*_*e*_ of the zoononoses (x-axis), the selective advantage *σ* of a mutation (Panel A) and the mutation rate *µ* (Panel B). *P*_*E*_ increases very steeply (the y-axis is on a log-scale) with increases in *R*_*e*_, and falls rapidly as *σ* and *µ* decrease. Parameters: *µ* = 10^−3^ in panel A and *σ* = 0.1 in panel B. The calculations for the probability of evolutionary emergence (defined as reaching an *R*_0_ ≥ 3 (really this should be *R*_*e*_ ≥ 3)) are described in the SI.

### Evaluating pandemic potential (*P*_*P*_)

We define the pandemic potential, *P*_*P*_ as the probability of a successful emergence event per unit time. The dependence of *P*_*P*_ on the probability of emergence per spillover, *P*_*E*_ and the spillover rate *S* is given by

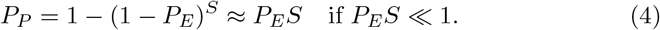

This is intuitive – when *P*_*E*_ is small the probability of a pandemic scales linearly with the number of spillover events and saturates as the probability of spillovers causing a pandemic approaches one. We first consider how *P*_*P*_ depends on the rate of spillovers (*S*) and the transmissibility of the pathogen in the human population (*R*_*e*_), and then apply this understanding to explore how *P*_*P*_ for mpox increases as population immunity to smallpox wanes.

The heatmap in Fig 4 shows *P*_*P*_ on a log scale, and we find that there is a very strong dependence on *R*_*e*_, particularly as *R*_*e*_ is slightly less than 1. In this regime *P*_*P*_ increases exponentially with linear increases in *R*_*e*_, consistent with the plots in Fig 3. In contrast *P*_*E*_ shows a much weaker, linear dependence with *S*.

**Figure 4:**
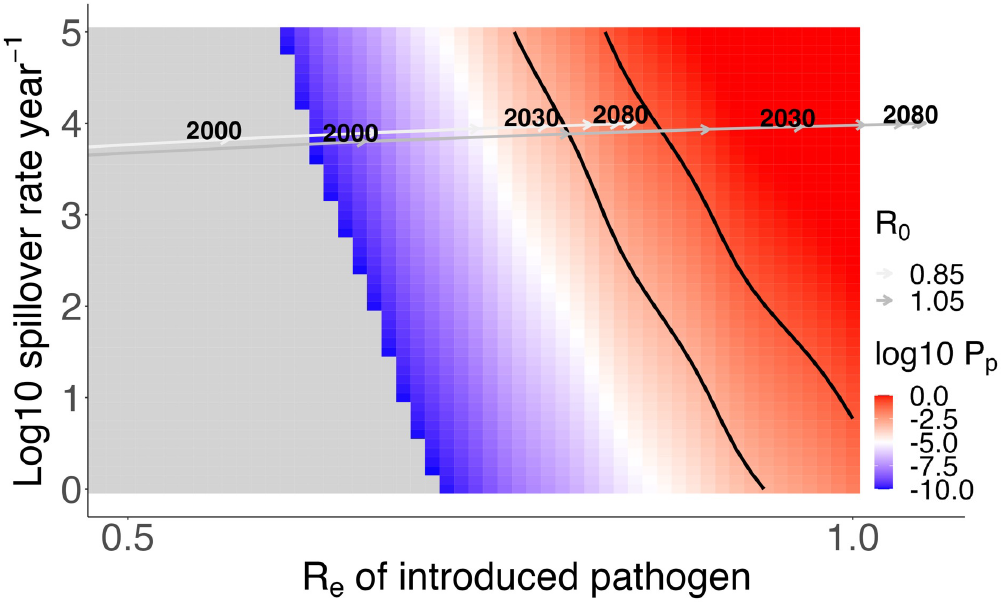
Pandemic potential *P*_*P*_ increases largely due to *R*_*e*_. We plot how *P*_*P*_ depends on *R*_*e*_ (x-axis) and *S* (y-axis). The black lines demarcate the region where 0.1 *< P*_*P*_ *<* 0.001 indicating the range where evolutionary emergence is likely to pose a problem. The gray lines with arrows show two example trajectories for how waning of population immunity to smallpox (taken from Fig 2) can result in a large increase in *P*_*P*_. These trajectories move from a region where emergence is unlikely to one where it is very likely, and this occurs largely due to the increase in *R*_*e*_ rather than *S* as population immunity to smallpox declines. Parameters: *s*_0_ = 10^4^, *σ* = 0.1 and *µ* = 10^−3^.

There is a narrow range of *R*_*e*_ for which the pandemic potential falls between and 0.001 per year, which corresponds to average times of emergence of 10 and 1000 years (black lines in Fig 4.) This is a relevant timeframe where evolutionary emergence poses a problem. If the time to emergence is shorter than 10 years than we might expect the pathogen to already have emerged, and if the time to emergence is greater than 1000 years disease emergence is unlikely to be of concern or even interest of policymakers or politicians. There is only a narrow band of *R*_*e*_ and *S* where 0.1 *< P*_*P*_ *<* 0.001, indicating few pathogens are likely to fall in this range.

The case is somewhat different for mpox. This is because with waning immunity to smallpox the *R*_*e*_ for mpox is almost certain to go from a value much less than one (about 0.3 in the 1980s) to a value around or slightly greater than one as the fraction of individuals with immunity to smallpox falls to close to zero in the next decade or two. This is indicated by the grey trajectories in Fig 4 which indicate how *P*_*P*_ increases as immunity to smallpox wanes. We use the same demographic model as Fig 2 to estimate the loss in the fraction of the population immune in the DRC through time. We choose to begin at the *R*_*e*_ = 0.3 from the estimate in the early 1980’s and show how *P*_*P*_ changes with time thereafter. We do so for a range of *R*_0_ of mpox (from 0.85 to 1.05) and note that varying *S*_0_ results in a nearly vertical translation of the trajectories. We see that the change in the *P*_*P*_ due to waning immunity is largely driven by increases in *R*_*e*_ rather than the change in *S*. In the SI we show that this result is robust to different *µ* and *σ*, as well as the addition of rare pre-adapted variants of the pathogen in the zoonotic reservoir.

### Interventions to prevent emergence

Interventions to reduce the pandemic potential, *P*_*P*_ can do so by reducing *R*_*e*_ (the transmissibility of the pathogen) or reducing *S* (the rate of spillovers) – the parameters *µ* and *σ* are properties of the pathogen-host system and are not under our control. We now consider the effect of interventions on reducing *P*_*P*_ in a regime where the *R*_*e*_ of mpox is around 1 – which is relevant at the current time and near future.

Immunization can reduce both *S* and *R*_*e*_ and as described earlier the main contribution to reducing *P*_*P*_ arises as a consequence of the reduction in *R*_*e*_. Another way of reducing *R*_*e*_ is by building accessible and efficient public-health and clinical healthcare systems. These could incorporate rapid testing, appropriate treatment reducing the duration of infection as well as quarantine measures as applicable.

A final way of reducing *P*_*P*_ and emergence is by reducing the ecological overlap between the zoonotic reservoir and human hosts which results in fewer spillovers (lower *S*) [30]. Our simulations suggest this is likely to result in only a modest reduction in *P*_*P*_, compared with the much larger reductions that would be obtained by interventions such as vaccination that decrease *R*_*e*_. However this does not take into account the costs to achieve similar fold reductions in these parameters, the potential for the reductions to cover multiple pathogens, or have other beneficial effects.

#### Costs of delaying intervention

It is widely held that prevention is better than a cure and this is true for the case of mpox, as we illustrate in Fig 5. In its simplest form prevention might occur by ensuring *R*_*e*_ (or more accurately combination of *R*_*e*_ and *S*) is kept below a critical level that corresponds to an acceptably low probability for evolutionary emergence as depicted in Fig 5. We see that for a wide range of parameters this requires keeping *R*_*e*_ to a level slightly below one (the level depends on *σ*, and *µ*). This might entail, for example a 20% reduction in transmission (if for example it requires reducing *R*_0_ of the zoonotic infection from 1 to 0.8). In contrast if mpox has had time to evolve its transmissibility to a level commensurate with smallpox which has an *R*_0_ about 3 then pathogen elimination requires preventing 1 − 1*/R*_0_ = 2*/*3 or 66% of all transmissions. Consequently, a much lower level of intervention such as vaccination is needed to prevent a pandemic rather than control it after it has occurred, and the pathogen has adapted to transmit in the human population as illustrated in Fig 5.

**Figure 5:**
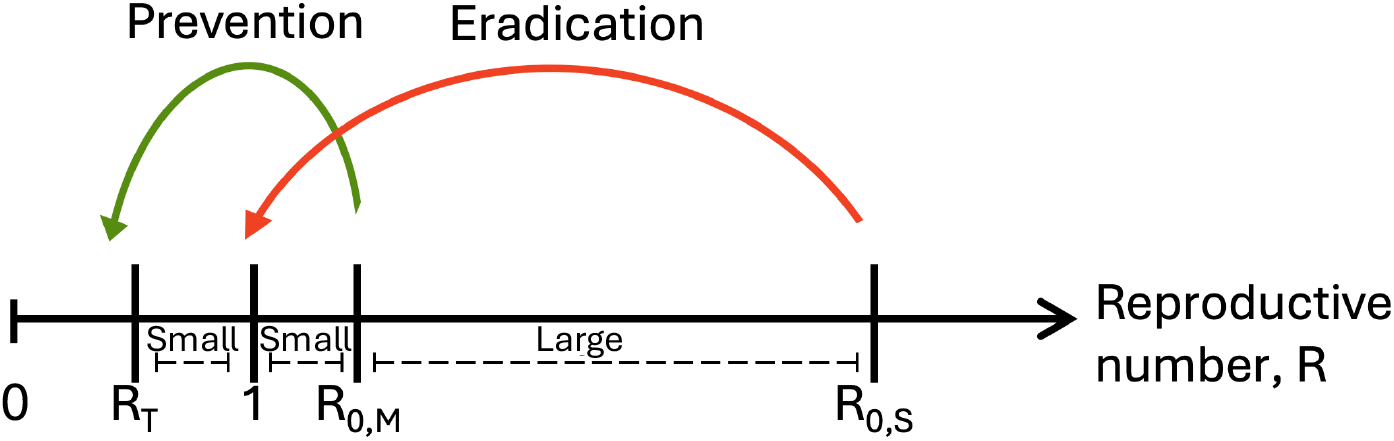
Eradication requires a greater intervention than prevention. The magnitude of the response required to control disease after its evolution to high transmissibility (*R*_0,*S*_) equals (1 − 1*/R*_0,*S*_). This is significantly greater than the magnitude required to prevent the emergence of the disease when its reproductive number is still small (*R*_0,*M*_), which equals (1 − *R*_*T*_ */R*_0,*M*_). *R*_*T*_ is the *R*_*e*_ for which *P*_*P*_ is sufficiently low.

## Discussion

Prior to the eradication of smallpox, mpox was not a problem. This was because cross-immunity arising from smallpox infection or vaccination largely prevented zoonotic spillovers as well as human-human transmission of mpox. The end of smallpox circulation and vaccination in 1980 has led to concomitant waning of immunity to mpox and an increase in both the number of zoonotic cases and the human-human transmission of mpox. We have used mathematical models to explore the interplay between waning immunity and evolutionary adaptation and applied this theory to the potential emergence of an evolved and highly transmissible mpox virus, and note its more general applicability to the emergence of zoonotic pathogens following the eradication of related endemic diseases.

Evolution is an inherently stochastic process that is naturally described by a branching process. We extended a previous model of the evolution of a pathogen during stuttering chains of transmission after spillover in three ways. First we incorporated changes (an increase) in the effective reproductive number (*R*_*e*_) for a single pathogen (mpox) as cross-immunity waned. Second, we integrated the increase in *R*_*e*_ with a decreases in the number of mutations needed for the virus transmission to become supercritical. Finally we allowed evolution continue even after *R*_*e*_ of the virus in the human population was greater than one.

Our analysis showed that as *R*_*e*_ approaches one there is a very rapid rise in the probability of evolutionary emergence, and this steep relationship is exacerbated by the decline in the number of mutations required as *R*_*e*_ approaches one. The rapid change in the probability of emergence with reproductive number suggests that there is only a small range of *R*_*e*_ for which emergence is likely to be a concern – if the probability of emergence is too small then it is not likely to occur in a time-frame of relevance (say, the next 100+ years), and if it is very high then the pathogen will have emerged or there will be little to prevent it emerging unless drastic measures are taken to reduce its reproductive number. The waning of immunity to mpox will result in its reproductive number increasing from well below one in the 1980’s (estimated at around 0.3) to close to or likely above one in the next decade when the population has almost no immunity to smallpox. Indeed, recent outbreaks of mpox have led to sustained human-human transmission [40]. Thus mpox may well be passing through the regime where emergence is likely now or in the near future.

Interventions to prevent the evolutionary emergence of mpox can do so by reducing the frequency of spillover infections (*S*) or the extent of transmission in the human population (*R*). Our results show that the latter (decreasing *R*_*e*_) is much more effective effective than the former (decreasing *S*). Ecological interventions that reduce contact between the zoonotic reservoir and humans can certainly reduce the number of zoonotic infections but unless the number of spillover events is very low our results suggest that this is unlikely to have a substantial effect on the probability of evolutionary emergence. Other interventions such as improvements in surveillance and universal access healthcare can reduce the transmission of emergent pathogens such as mpox in humans and have the advantage of doing so not just for mpox, but also for other zoonoses and would benefit many aspects of the health of communities.

Vaccination can decrease emergence by reducing both the spillover rate and transmission of a disease, and our study suggests that its effect is largely from the extent of reduction in transmission. Vaccines can work in multiple ways – they can reduce the susceptibility of individuals to infection, and they can reduce the infectivity and the extent of pathology in those that do get infected. Preventing the evolutionary emergence of mpox by vaccination would benefit from the design of a vaccine that confers broad orthopox immunity potentially at the expense of immunity to smallpox and would also require a focus on reducing transmission (by a combination of reducing susceptibility and infectivity) rather than pathology. Indeed a vaccine that only reduces pathology might increase the probability of emergence due to transmission from asymptomatic infections. It may even be that permitting some symptoms can allow for the better detection of spillover infections and facilitate measures that reduce transmission [14].

Perhaps the most salient result is the observation that the level of vaccination or other intervention required to prevent emergence is modest compared to that needed for control and eradication after it has adapted to high transmissibility in the human population post-emergence. Due to the persistence of the pathogen in zoonotic reservoirs, however, this low-level vaccination needs to be ongoing and introduces an interesting difficulty with the counterfactual. To what extent can we know the true necessity of prevention efforts if the disease never emerges? We leave this question to the reader as a philosophical exercise.

### The epidemiology of mpox indicates limitations of our understanding

The framework we have described applies to mpox, the only human disease which we have eradicated. As might be expected and consistent with our model mpox has become an increasing problem since vaccination against smallpox was discontinued, and the fraction of individuals with smallpox immunity declined. In recent years *R*_*e*_ of mpox has reached around 1 in groups where close contact, typically during sex, facilitates its transmission. This has resulted in the disease persisting in these groups, even in the absence of ongoing zoonotic introductions [13]. Fortunately the reproductive number of both clade 1 and 2 strains in these groups is only slightly greater than one, allowing modest public health interventions and targeted vaccination to contain but not eliminate these outbreaks. Over time the waning of residual population immunity may allow a further increase in the *R*_*e*_ of mpox allowing it to sustain transmission in a larger fraction of the population. The extent to which this occurs is not likely to be large as at present only a small fraction of the population (those born before 1980) has been vaccinated against smallpox and furthermore their immunity has likely waned considerably over the past decades. However the ongoing human to human transmission in the high risk groups in which it is currently endemic raises the probability of it adapting to increase transmission between humans. This would result in mpox being able to spread in more communities, and necessitate much more widespread intervention for its control. We now discuss some of the complexities associated with mpox epidemiology and evolution that limit our ability to make quantitative predictions of its evolutionary trajectory.

Mpox could adapt by increasing its ability to replicate within individuals, evade the immune response or transmit via a respiratory route in addition to the very close contact needed for transmission of the currently circulating strains [10, 18]. A major limitation is our lack of knowledge of the mutational and adaptive landscapes that underlie these changes, and research in this area is necessarily limited by the safety of experimental evolution of mammalian-infecting viruses [23]. It is likely that virus adaptation requires multiple mutations and this accounts for why we have not yet seen variants with much higher fitness despite clades 1 and 2 having infected over 50,000 and 100,000 individuals respectively [13]. This was noted for SARS-CoV-2 where it took several months and millions of infections before more transmissible delta and omicron variants emerged – and these viruses had dozens of mutations [38]. Circulating mpox variants contain many more mutations than expected, but this is most likely due to editing by human APOBEC3 enzyme, which are indicative of sustained human to human transmission as opposed to multiple zoonotic introductions or adaptation [29, 36]. Whether some of these mutations increase mpox fitness in the human host remains to be seen. We also note that the branching process calculations which estimate the probability of evolutionary emergence do not take into account the interval between the time at which the successful zoonotic event occurs, the time at which the evolved variant is generated (which could involve a long chain of transmission events), and the time at which it has reached a sufficiently high frequency to be recognized as a public health concern. Given that mpox appears to have a relatively long serial interval, the time delays of associated with these two processes could be substantial [26, 31].

We also need to consider heterogeneity in transmission events which has been the subject of considerable interest [25], and is a potential direction for extensions of the current model. The dynamics of infection and immunity of mpox in high risk groups is of particular importance and it may be very different from that of other pathogens. The persistent nature of HIV and many STD’s results in highly connected nodes of sexual networks contributing substantially to the spread of these STD’s. In the case of acute infections such as mpox infections of individuals in highly connected nodes would result in surviving individuals being immune and this would potentially limit further spread. Regardless, as population level immunity is lost we would expect to see the first signs of increased transmission in outbreaks among high transmission groups.

Another problem is that the spread of an emerging pathogen can result in changes in the contact patterns of humans and modify its spread, and the extent of these changes can vary dramatically between different locations as was seen following the initial emergence of SARS-CoV-2 where some countries could prevent successful introductions (e.g. New Zealand), a few could introduce sufficiently severe lock-downs that resulted in local eradication (e.g. China) and most could at best flatten or reduce the number of cases prior to development of a vaccine [21, 22].

In conclusion our limited understanding of disease emergence arises both from complexities in the evolutionary and epidemiological processes involved and the paucity of examples of pathogens that have successfully emerged into the human population recently enough to allow detailed analysis. Indeed, mpox is currently the strongest example of a zoonotic disease for which the loss of cross-protective herd immunity might facilitate its emergence. Still, while the actual eradication of smallpox required a far more nuanced strategy that included surveillance followed by ring vaccination, the simple formulation of a critical threshold for vaccination *f >* 1 − 1*/R*_0_ provided the rationale for the eradication of smallpox. Similarly the simple analysis we describe highlights the potential risk of evolutionary emergence associated with disease eradication and the subsequent waning of immunity to related zoonoses. The considerations outlined in this paper might inform us of how early and ongoing intervention might be needed following the eradication of pathogens such as smallpox.

