## Supplemental Information for "Preventing Disease Emergence Following Eradication: Application to Mpox"

### 1 Multi-type branching process formulation

Our model for the adaptive evolution of a zoonotic pathogen after spillover into humans is based on a previously developed multi-type branching process model of emerging infectious diseases [1]. The multi-type branching process formulation allows for an easy derivation for the probability of emergence.

Consider the case where a pathogen must accumulate a number of mutations after spillover into the human population to attain $R_0 > 1$. Given $m$ variants from the wildtype to the fully-evolved mutant with $m-1$ mutations required to attain $R_0 > 1$, the $m$-type branching process can be described by $m$ probability generating functions (PGFs):

$$f_i(s_1, s_2, ..., s_m) = \sum_{j_1, j_2, ..., j_m = 0}^{\infty} p_i(j_1, j_2, ..., j_m) s_1^{j_1} s_2^{j_2} ... s_m^{j_m}, \ i \in \{1, 2, ..., m\}$$

where $p_i(j_1, j_2, ..., j_n, ..., j_m)$ is the probability that an infection of type $i$ gives rise to $j_n$ secondary infections of type $n$.

The probability of extinction $q_i$ of a transmission chain initiated by a single infection with a variant of type $i$ is given by the least fixed-point solution to: $f_i(q_1, q_2, ...q_m) = q_i, \ i \in \{1, 2, ..., m\}$.[1]

We define a model in which there exists $m$ different variants of the pathogen, with one wildtype and $m-1$ mutants. We denote the basic reproductive number of the wildtype pathogen as $R_0^{(1)}$, the basic reproductive numbers of the $m-2$ intermediate mutants as $R_0^{(2)}, ..., R_0^{(m-1)}$, and the basic reproductive number of the fully-evolved mutant as $R_0^{(m)}$.

We assume the following about mutation and the spread of infections:

- the mutation rate, $\mu$, of all the variants is the same

[1] Note that this is a special case of the generalized expression for the probability that a process initiated with $r_1, r_2, ..., r_n$ chains of infections of type $1, 2, ..., m$ goes extinct, which is given by the product: $q_1^{r_1} q_2^{r_2} ... q_m^{r_m}$.

- only single variants occur, i.e the $it$th variant can only mutate into an $i+1$ mutant
- the total number of secondary infections arising from an individual infeected with variant $i$ is Poisson-distributed with mean $R_0^{(i)}$
- the number of secondary infections of type $i$ arising from an individual infected with variant $i$ is Poisson-distributed with mean $(1-\mu)R_0^{(i)}$
- the number of secondary infections of type $i+1$ arising from an individual infected with variant $i$ is Poisson-distributed with mean $\mu R_0^{(i)}$

These assumptions define the PGFs $f_1, f_2, ..., f_m$, given by:

$$f_i(s_1, s_2, ..., s_m) = \exp[(1-\mu)R_0^i(s_i-1)]\exp[\mu R_0^i(s_{i+1}-1)],\ i \in \{1, 2, ..., m-1\}$$

$$f_m(s_1, s_2, ..., s_m) = \exp[R_0^m(s_m-1)]$$

The probability of emergence can be calculated numerically from the PGFs according to the fixed point equation above.

Our model makes 2 important departures from the previous model:

1. The fitness of intermediate variants increases by some proportional amount, $\sigma$;
2. Evolution continues to increase the reproductive number after the first $R > 1$ variant.

We alter the framework presented in [1] according to the following.

Consider the case of adaptive evolution during transmission. After the spillover infection, mutations are acquired during subsequent transmission events, each increasing the $R$ of infections by some proportion, $\sigma$, until a variant emerges with an $R_0^{(m)}$ exceeding some $R_0^{(m')}$.

The number of mutations $m-1$ in the lineage from the wild-type (1) to the fully-adapted type ($m$) thus depends on the reproductive number of the wildtype $R_0^{(1)}$, the minimum reproductive number of the fully-adapted type $R_0^{(m')}$, and the proportional increase in $R_0^{(i)}$ to $R_0^{(i+1)}$ with mutation and is given by:

$$m-1 = \frac{log(R_0^{(1)} - log(R_0^{(m')})}{log(1+\sigma)}$$

Note that $R_0^{(m)}$ is the first $R_0$ that is greater than the minimum $R_0^{(m')}$.

To calculate the probability that the lineage does not go extinct before exceeding $R_0^{(m')}$, we replace the final $R_0^{(m)}$ with some R arbitrarily high such that the contribution of extinction of the final type is negligibly small (once the final type arises the process will never go extinct, so extinction probability is determined by intermediates.)

### 2 Strong dependence of P(Emergence) on Re is Robust to $\mu$ and $\sigma$

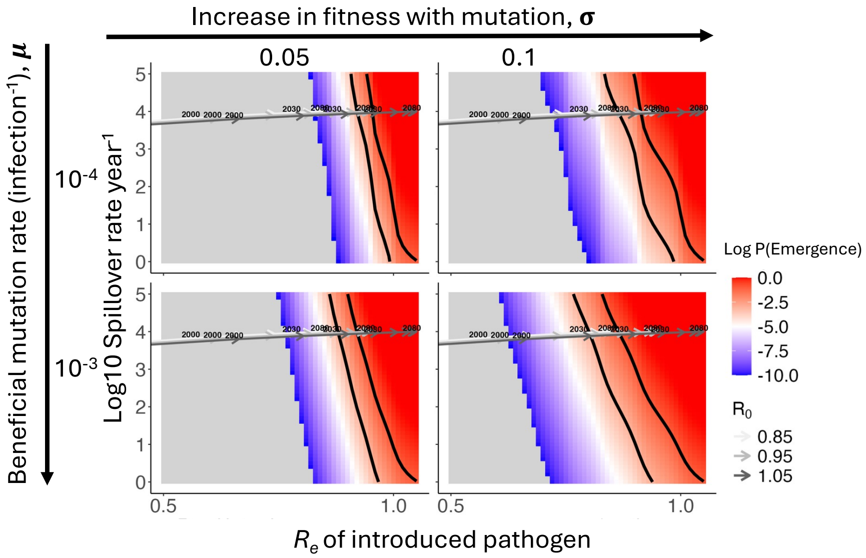


Figure 1: **Effect of $\sigma$ and $\mu$ on the trajectory of pandemic potential P(emergence) as immunity wanes.** The trajectories of S0 and Re do not change as mu and sigma vary, but the time spent in the relavant regime does. The strong dependence of the change in P(emergence) on Re rather than S is robust to changes in the pathogen characteristics $\mu$ and $\sigma$.

### 3 Strong dependence of $P_p$ on Re is also characteristic of a pre-adaptation pathway.

We now consider pre-adapted variants of a pathogen that circulate in the zoonotic reservoir. Such as pre-adapted pathogen variant might for example require one fewer mutation for successful adaptation to transmission between humans. The rate of spillover of this variants into humans would be proportional to their frequency in the zoonotic reservoir which is likely to be low and reflected by a decrease in S (due to mutation-selection-drift balance). Changes in pandemic potential are still largely driven by increase in $R_e$ rather than $S$ along this pathway, with only a slightly reduced effect owed to the decrease in the number of mutations required for emergence.

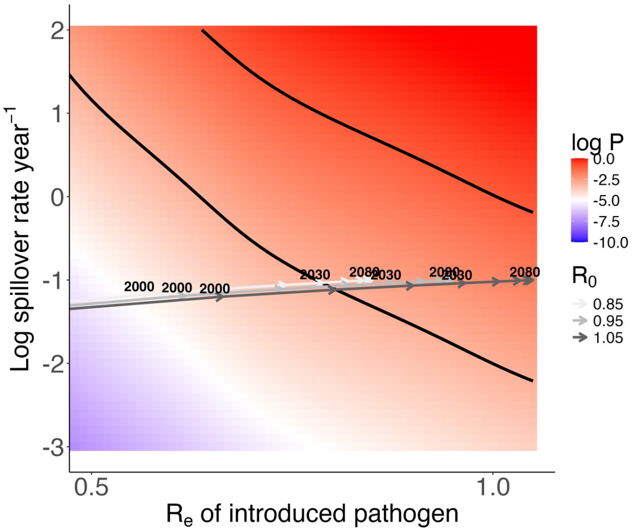


Figure 2: **In a pre-adapted pathway, variants with high $R_e$ and that are few infections away from emergence (due to large $\sigma$ of $mu$ may circulate endemically in the animal reservoir.** Assuming that the mutations conferring pre-adaptation to humans are not so deleterious that the variants cannot persist in the animal reservoir, they may circulate at low frequency (mutation-selection-drift balance.) The frequency of spillovers, $S$, would be proportional to the frequency in the zoonotic reservoir, and thus $S$ would be in a significantly smaller regime. The dependence of pandemic potential on $R_e$ rather than $S$ is maintained in this regime, and thus interventions targeting transmission may have the largest effect on preventing emergence ($S$ of all variants may be high and hard to reduce on a log-scale).
